# Segmental Isotope Labelling of the Prion Protein: Identification of a Key Residue for Copper-Mediated Interdomain Structure

**DOI:** 10.1101/2025.05.01.651769

**Authors:** Francesca A. Pavlovici, Kevin Singewald, Samuel Kaplan, Eefei Chen, Glenn L. Millhauser

## Abstract

The cellular prion protein is composed of two domains: a disordered N-terminal toxic effector domain and a three-helix C-terminal regulatory domain. Copper is thought to form a bridge between these two domains, inhibiting the protein’s inherent neurotoxicity. However, the molecular details of how copper interacts with the C-terminal regulatory surface are unclear. To assess the potential role of conserved C-terminal His residues in copper coordination, we applied sortase-mediated ligation to create an expressed, murine prion protein with segmental ^15^N-labeling of the N-terminal domain. Pulsed EPR methods applied to a 1:1 protein:copper complex revealed both ^14^N and ^15^N couplings, consistent with simultaneous coordination of the two protein’s domains to the copper center. Mutagenesis studies localized C-terminal copper coordination to His176, present on the second α-helix. The cumulative EPR results reveal a copper coordination environment composed of three His residues from the protein’s N-terminal domain, along with His176. The feasibility of these findings was tested with AlphaFold 3 simulations. These results further refine the molecular details of the prion protein’s autoregulation, emphasizing the critical role of its copper cofactor. Moreover, this interdisciplinary work demonstrates how sortase-mediated ligation combined with pulsed EPR sensitive to distinct nuclear spin systems provides a new strategy for assessing metal ion binding to proteins.

## Introduction

Prion diseases are transmissible, fatal neurodegenerative illnesses in humans and both domestic and wildlife animals.^1,2^ These diseases arise from the misfolding and subsequent aggregation of the endogenous, cellular prion protein (PrP^C^) into its pathological isoform, referred to as the scrapie prion protein (PrP^Sc^).^1,3^ PrP^C^ is a glycosylphosphatidylinositol (GPI) anchored, glycoprotein expressed throughout the body, but primarily in the central nervous system.^4–7^ PrP^C^-null mice, when inoculated with exogenous PrP^Sc^, fail to develop prion pathology or neurodegeneration.^8–11^ This suggests that the presence of PrP^Sc^ alone is not sufficient to promote neuron dysfunction. Instead, the current paradigm posits that PrP^Sc^ acts by binding to GPI-anchored PrP^C^, driving neurotoxic processes at the cellular membrane.^12^ In further support of this concept, mice expressing “anchorless” PrP^C^, when inoculated with PrP^Sc^, show widespread protein aggregation but are slow to show the neurotoxic effects associated with prion disease.^13^ Finally, electrophysiological experiments with cells expressing select mutants of monomeric PrP^C^ reveal spontaneous cationic transmembrane currents, suggesting that neither aggregation nor the presence of PrP^Sc^ are required for PrP^C^ neurotoxicity.^14,15^

Murine PrP^C^ is composed of a flexible N-terminal, toxic effector domain (residues 23-125), and a globular C-terminal regulatory domain (residues 126-230) (**Figures 1A and 1B**).^15,16^ Within the N-terminal domain is the octarepeat (OR) segment, composed of tandem repeats of the eight-residue sequence PHGGXWGQ (where X is Gly in repeats one and four and Ser in repeats two and three in the mouse sequence) (**Figure 1A**).^17,18^ Also within the N-terminal domain is the Central Region (CR) linker (residues 105-125), the deletion of which drives profound neurotoxicity in both transgenic mice and in immortal cell lines.^15,19,20^ The N-terminal toxic effector domain of PrP can interact with the lipid bilayer or with other membrane proteins to cause several toxic activities, including membrane leakage, spontaneous ionic currents, and neuronal death.^15,16,19,21,22^ Cu^2+^ coordinates simultaneously to His residues in the OR segment^17,23–26^ and a somewhat diffuse, negatively charged patch on the globular C-terminal domain (**Figure 1B**),^27–29^ suppressing these deleterious activities.^15^ This neuroprotective, tertiary fold is termed the Cu^2+^-driven *cis* interdomain interaction (hereafter referred to as the *cis* interaction). Consistent with the recognition of PrP^C^ as a copper binding protein, spatially resolved brain Cu^2+^ levels correlate with PrP^C^ expression.^30,31^

**Figure 1.**
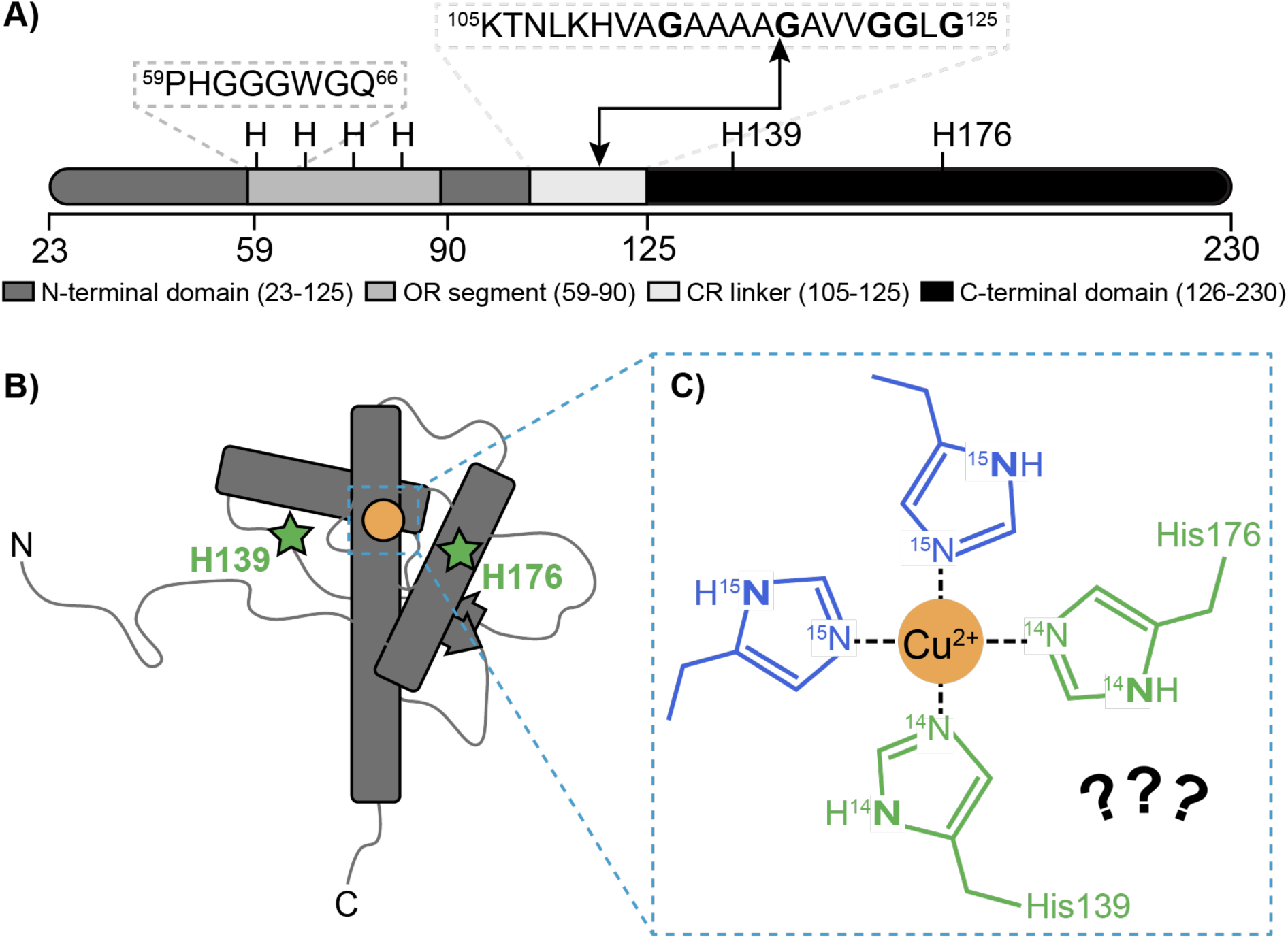
Copper is Proposed to Form a Bridge Between the Prion Protein’s Domains. **A)** Murine PrP^C^ is composed of two major domains, the N-terminal domain (residues 23-125) and the C-terminal domain (residues 126-230). Within the N-terminal domain are two sequence motifs, the octarepeat (OR) segment (residues 59-90) and the central region (CR, residues 105-125). The OR segment is comprised of eight amino acids, PHGGGWGQ, repeated four times. The black arrow points to the domain boundary (Ala117/Gly118) in the CR that creates the two precursors for the ligation reaction. Annotated are four His residues (His60, His69, His76, His84) in the OR and two (His139 and His176) in the C-terminal domain relevant to Cu^2+^ binding. **B)** The N-terminal domain is intrinsically disordered; the C-terminal domain is globular, composed of three α-helices (cylinders) and two β-strands (arrows), forming an antiparallel β-sheet. Previous work finds that the OR-bound Cu^2+^ ion (gold sphere) associates with a surface region of the C-terminal domain forming a bridge between the two major domains via His residues (green stars). **C)** The mechanism by which Cu^2+^ bridges between the two domains may involve coordination by domain specific His residues (OR His in blue, C-terminal His in green). The bolded nitrogen atoms are detectable by ESEEM and HYSCORE.

Our lab has sought to reveal the molecular details of the *cis* domain-domain interaction, facilitated by the copper bridge (**Figure 1C**).^27,29^ Despite the importance of this self-regulation mechanism, only limited information is known about the regulatory interface between the protein’s two domains. Our current model^27,32,33^ is based on distance measurements derived from double electron-electron resonance (DEER) EPR^34,35^ with spin labeled PrP^C^ and NMR experiments that identify contact surface residues within the C-terminal domain, as reflected by paramagnetic relaxation enhancement (PRE).^27,29,32^ The PRE experiments find that over 20 C-terminal residues are affected, including two C-terminal His residues at positions 139 and 176 that may serve as primary Cu^2+^ contact points through metal ion coordination. However, neither EPR nor NMR, as we have previously applied the methods, conclusively identifies C-terminal His coordination. For example, while the pulsed EPR method of electron spin echo envelope modulation (ESEEM)^36^ applied to Cu^2+^ centers is definitive for His coordination,^37–39^ the resulting signals cannot distinguish between His residues from the N- or C-terminal domains.^29^ Here, we further elucidate the molecular details of the *cis* interaction by developing an approach that combines the chemical biology of protein ligation along with newly developed pulsed EPR measurements to rigorously test for C-terminal His coordination to the Cu^2+^ center.

We recently demonstrated that a Cu^2+^ ion bound to ^14^N- or ^15^N-His residues gives distinct, fully resolvable signals in both ESEEM and the two-dimensional EPR method of Hyperfine Sublevel Correlation Spectroscopy (HYSCORE).^40,41^ In that previous work, one protein in a two-protein complex was uniformly ^15^N-labeled. By applying ESEEM and HYSCORE to the copper-stabilized protein complex, we not only observed simultaneous coordination of the two proteins to the copper center, but we also evaluated the number of His residues contributed by each. Here we apply a similar approach to assess the relative domain contributions within a single protein. Specifically, we used sortase-mediated ligation (SML)^42–47^ to produce PrP^C^ with ^15^N-labeling exclusively to the N-terminal toxic effector domain (**Figure 2A**). This ligation strategy applied to PrP^C^ retains ^14^N-His side chains at positions 139 and 176, which, if Cu^2+^-coordinated through the *cis* interaction, will be readily resolvable by ESEEM and HYSCORE. The findings were further developed and expanded using site-directed mutagenesis to test and refine the assignments.

**Figure 2.**
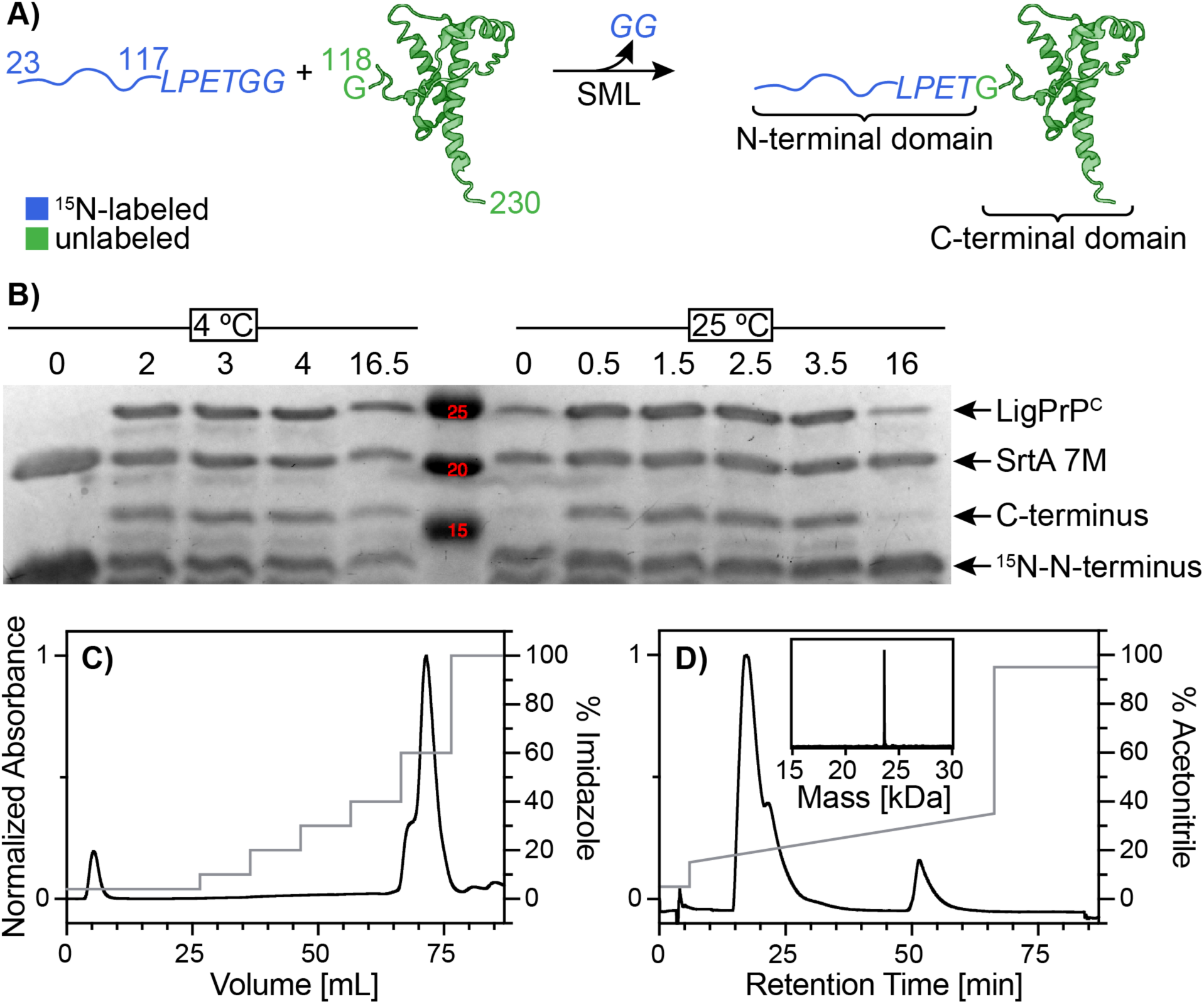
Sortase Mediated Ligation Applied to the Prion Protein. **A)** The LPXTG substrate is composed of PrP^C^ residues 23-117 appended with the sortase handle *LPETGG* (blue), expressed with uniform ^15^N labeling. The glycine nucleophile, PrP^C^ residues 118-230, possesses a native Gly at its N-terminus. Upon ligation, the Gly-Gly dipeptide is released. The final fusion product is native, murine PrP^C^ with an LPET insert (italicized) between Ala117 and Gly118. Created in BioRender. (2025) https://BioRender.com/p4nxprc **B)** Ligation reactions at 4 °C and room temperature had time points collected over the course of 16 hours. Analysis by SDS-PAGE gel found that ligation reactions at room temperature reach equilibrium after 30 minutes. The numbers (red) are the molecular weights in kDa of protein standards’ bands which are used as references to identify protein bands in the other lanes. **C)** Elution of reaction components were monitored by absorbance at 280 nm. An imidazole step gradient (gray, from 4 mM to 250 mM imidazole) purified away the unreacted C-terminal domain (volume 5 mL, 4% imidazole) and SrtA 7M (volume ∼85 mL, 100% imidazole). The major peak (volume starting at 65 mL, 60% imidazole) contained both excess ^15^N-N-terminal domain and LigPrP^C^. **D)** Final purification by HPLC monitored by absorbance at 280 nm with an acetonitrile linear gradient (gray, from 15% to 35% acetonitrile) isolated LigPrP^C^ (retention time starting at 50 minutes) from excess ^15^N-N-terminal domain (retention time 15 minutes). The inset reveals the peak at 50 minutes gave a mass of 23,646 Daltons.

## Results and Discussion

### Creation and Characterization of LigPrP^C^

Our goal was to use Sortase A heptamutant (SrtA 7M)^48^ to produce PrP^C^ with a uniformly ^15^N-labelled N-terminal domain (**Figure 2A**). To select the best ligation point within the PrP^C^ sequence, we considered several critical factors. First, we desired a sequence that left only the folded C-terminal domain with ^14^N-His residues. Next, to facilitate reactivity, we considered the solubility of the two reacting fragments, as well as exposure and flexibility of the ligating termini.^49–52^ Finally, we sought to minimize changes in the fundamental PrP^C^ sequence. To achieve these goals simultaneously, we targeted the reaction to the position 117-118 (**Figure 1A**). Residue 118 is natively a Gly residue and meets the requirement for SrtA reactivity of the C-terminal domain. The reacting C-terminal residues of the N-terminal domain’s “ligation handle” typically require the sequence LPXTG.^53^ We selected a Glu residue for position X, which has been found to facilitate the reaction and enhance solubility.^54^ We also extended to an additional Gly residue to further facilitate ligation reactivity (**Figure 2A**).^50^ The resulting LPETGG handle was appended to residue 117 of the N-terminal domain. The resulting reaction is shown in Figure 2A and results in the native PrP^C^ sequence with a tetrapeptide insert of LPET following residue 117. We refer to this construct as LigPrP^C^.

To optimize the SML, we varied reactant concentrations,^51,55^ pH,^51^ temperature^56,57^ and reaction time^50,56^ (**Figure S1**). Ultimately, we found that a 10:5:1 ratio for N-term, C-term, and SrtA 7M, respectively, at 100 µM concentration of N-term provided the greatest product yield. We carried out the SML at room temperature and pH 7.4 (not at the canonical pH 7.5) to maintain solubility of the C-terminal domain, while meeting the pH requirements for SrtA activity (**Figure 2B**). Back reaction of SML was minimized by dialyzing out the Gly-Gly leaving group through the course of the reaction.^49,50^

Because all protein components of the mixture are rich with His residues, standard Ni^2+^ gravity column purification was not sufficient to isolate LigPrP^C^. To overcome this, we employed Ni^2+^-immobilized metal affinity chromatography (Ni^2+^-IMAC), which facilitates the use of imidazole gradients to purify away SrtA and unreacted PrP^C^ C-term (**Figure 2C**). Following this, HPLC was used to capture the LigPrP^C^, separating this product from the excess N-term (**Figure 2D**). HPLC revealed a single peak at retention time of 50 minutes, giving a mass of 23,646 Da, corresponding to the molecular weight of LigPrP^C^ (**inset of Figure 2D**).^52^ The final yield of pure LigPrP^C^ is approximately 20% of the C-terminal PrP^C^ construct, ultimately providing approximately 500 micrograms of LigPrP^C^, fully sufficient for EPR studies.

To establish the utility of LigPrP^C^, we performed both CD and EPR control experiments shown in Figure 3. For helical proteins, such as the C-terminal domain of PrP^C^, CD spectroscopy should give negative dichroic bands at 208 and 222 nm.^58^ These transitions are clearly observed for both PrP^C^ WT and LigPrP^C^ (**Figure 3A**) with signal intensities matching those previously published.^59^ (Note that PrP^C^ folds to its helical form in the absence of copper. The spectra in **Figure 3A** were obtained with one equivalent of Cu^2+^, thereby matching the samples prepared for EPR.) The similarity of the magnitudes of the respective CD spectra suggests that LigPrP^C^ adopts the global fold of PrP^C^. We conclude that the LPET insertion, which is outside of the globular domain, has negligible effect on the PrP^C^ three-dimensional structure.

**Figure 3.**
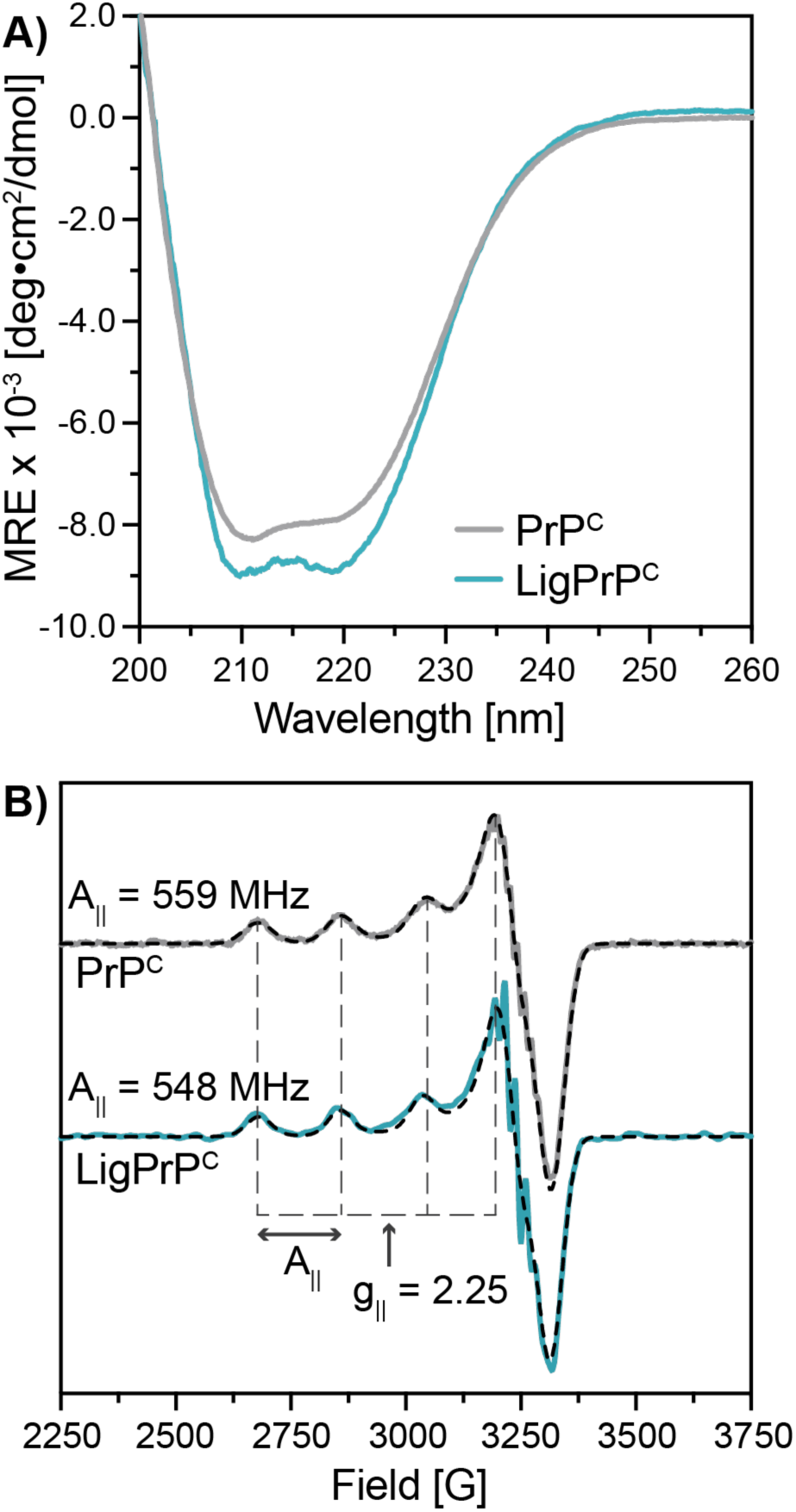
LigPrP^C^ is a Surrogate for PrP^C^. **A)** CD spectra of LigPrP^C^ (cyan) and PrP^C^ (gray) have negative bands at 208 and 222 nm, indicating similar α-helical content. **B)** CW EPR spectra of LigPrP^C^ and PrP^C^ with one equivalent of Cu^2+^. The CW spectra are fit using EasySpin (dashed lines). Values of g_||_ and A_||_ are indicated in the figure and are consistent with a 4N coordination environment.

Next, we performed X-band continuous wave (CW)-EPR experiments to assess the ligands involved in Cu^2+^ coordination. In the parallel region of copper EPR spectra, g_||_ and A_||_ values are sensitive to the nature of the coordinating atoms and the net charge of the copper center.^60^ We previously reported these values for PrP^C^ bound with one equivalent of Cu^2+^ (Component 3 coordination) and found g_||_ = 2.25 and A_||_ = 576 MHz, consistent with 3N1O or 4N coordination, arising from multiple His residues.^17^ CW-EPR spectra obtained from WT PrP^C^ (with no ^15^N-labeling) and LigPrP^C^ are nearly superimposable with each other in the parallel region and, when fit with EasySpin,^61^ yield g_||_ and A_||_ for LigPrP^C^ of 2.25 and 548 MHz, within experimental error of our previously published values (**Figure 3B**). These data demonstrate that the subtle changes in sequence arising from ligation do not perturb Cu^2+^ coordination. Additionally, the perpendicular region of the LigPrP^C^ spectrum at 3250 Gauss displays sharp superhyperfine peaks, characteristic of coupling from directly coordinated ^15^N-nuclei. CW-EPR spectra for all PrP^C^ constructs tested in this study were collected (**Figure S2**).

### Pulsed EPR Applied to LigPrP^C^ as a Test for C-terminal His Coordination

Our primary tools for characterizing Cu^2+^ coordination in LigPrP^C^ are ESEEM and HYSCORE. These pulsed EPR experiments detect energy transitions influenced by electron-nucleus coupling interactions. Contributions to the ESEEM transition energies include the nuclear Zeeman (NZ), hyperfine (HF), and nuclear quadrupolar interactions (NQI) (when I ≥ 1).^36,62^ In the case of copper proteins, these methods are diagnostic of His coordination through magnetic coupling to the remote nitrogen of the imidazole ring (**Figure 1C**). Moreover, as we have recently shown, ESEEM spectra give distinct transitions for ^14^N-His and ^15^N-His, and also reflect the number of coupled nuclei for the latter.^41,63^ Figures 4A and 4B depict the energy diagrams for the nuclear spin environments of a single coupled ^14^N (I = 1) and one or two coupled ^15^N (I = ½) His residues. For ^14^N at X-band frequencies, the Zeeman and HF interactions in the high energy manifold are in near “exact cancellation,” leaving only the prominent NQI transitions (**Figure 4A**).^64^ Consequently, we anticipate three quadrupolar transitions from the high energy manifold peaks, ν_+_, ν_-_, ν_0_, and a fourth double quantum (DQ) transition in the low energy manifold, ν_dq_. When an electron interacts with a ^15^N-nucleus, spectra reflect the NZ and HF interaction, giving two transitions: ν_α_ and ν_β_ (**Figure 4B**).^65^ If the HF interaction is negligible, these two transitions converge into a single ν_nz_. Two equivalent ^15^N-nuclei simultaneously interacting with the Cu^2+^ electron may give an additional weak 2ν_α_ transition.^41^ From this analysis and based on our previous characterizations, we find that both ^14^N and ^15^N give energy transitions that act as fingerprints reflecting the isotopic composition of the coupled His residues to the copper center.^41^

**Figure 4.**
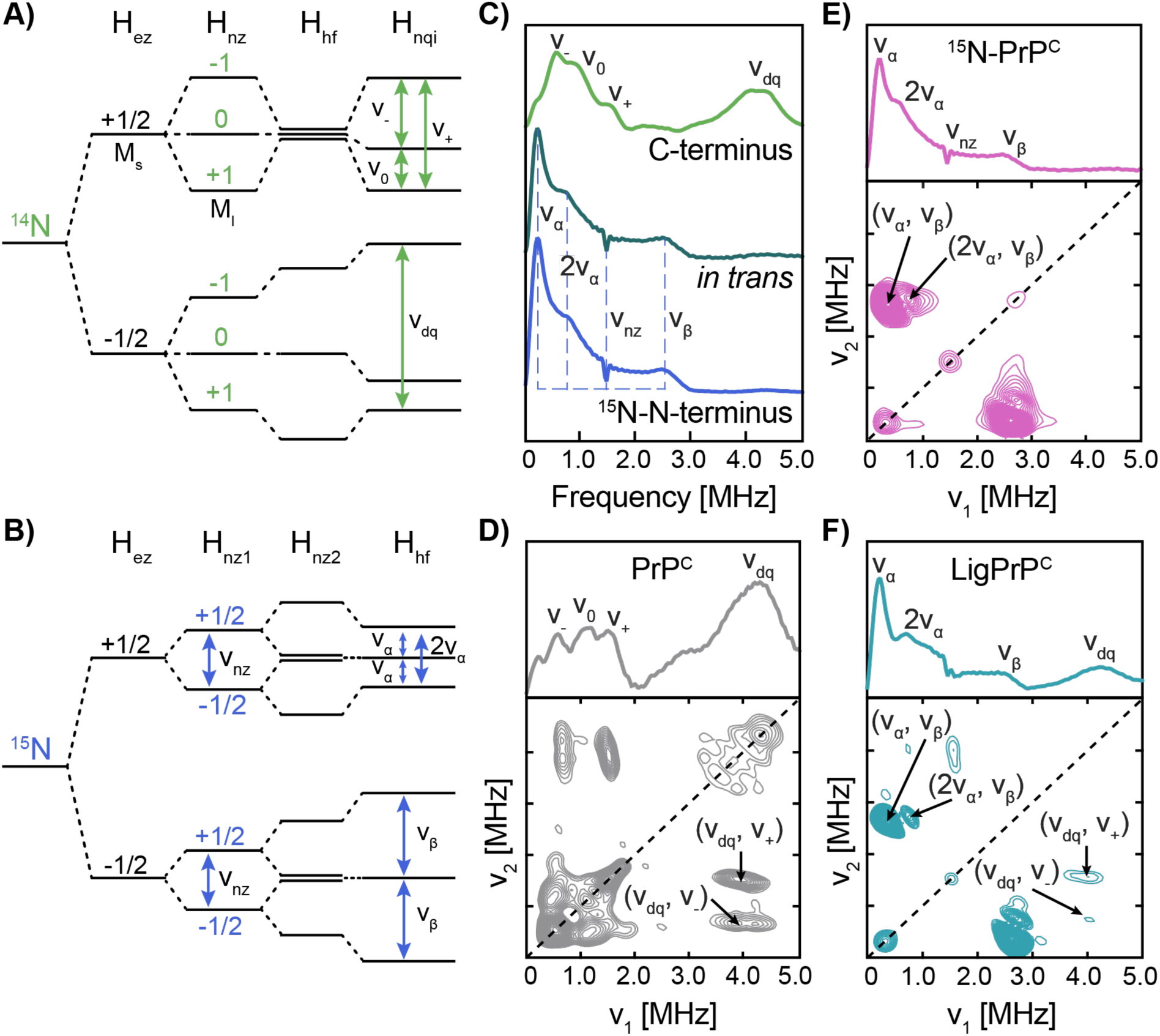
Pulsed EPR Experiments Reveal that in 1:1 Mixtures Cu^2+^ Simultaneously Coordinates to Both PrP^C^ Domains in LigPrP^C^. **A)** The energy transitions observed for a Cu^2+^ ion coupled to a ^14^N-nucleus (I = 1) are ν_-_, ν_0_, ν_+_, and ν_dq_. Direct NQI transitions are observed for the remote His imidazole nitrogen. **B)** When two ^15^N-nuclei (I = ½) couple to a Cu^2+^ ion, the observed energy transitions are ν_α_, 2ν_α_, ν_nz_, and ν_β_. **C)** ESEEM spectra of the unlabeled C-terminal domain (green) and the ^15^N-labeled N-terminal domain (blue). The ESEEM spectrum of *in trans* sample (1:1:1 ternary mixture) produces the four ^15^N energy transitions arising from the ^15^N-N-terminal domain, consistent with coordination by two or more His imidazoles. **D)** ESEEM and HYSCORE spectra of PrP^C^ with one equivalent of Cu^2+^ gives the NQI and DQ peaks and the ^14^N cross peaks of (ν_dq_, ν_-_) and (ν_dq_, ν_+_), respectively. **E)** ESEEM and HYSCORE spectra of ^15^N-PrP^C^ with one equivalent of Cu^2+^ gives the ^15^N-peaks of ν_α_, 2ν_α_, ν_nz_ and ν_β_ as well as the ^15^N cross peaks of (ν_α_, ν_β_) and (2ν_α_, ν_β_), respectively. **F)** The ESEEM spectrum of LigPrP^C^ with one equivalent of Cu^2+^ shows both the ^15^N-peaks of ν_α_, 2ν_α_, ν_β_ and the ^14^N-peak of ν_dq_, signals all of which are found as their corresponding cross peaks in the HYSCORE spectrum.

We first performed ESEEM on the ^15^N-N-terminal domain and the natural abundance C-terminal domain as separate peptides, each with one equivalent of Cu^2+^, to provide initial reference spectra (**Figure 4C**). The ESEEM spectrum of the ^15^N-N-terminal domain alone reveals transitions of ν_α_, 2ν_α_, ν_β_, and ν_nz_. Observation of the 2ν_α_ peak, as noted above, reflects multiple ^15^N-His coordination from the OR segment. The ESEEM spectrum of the C-terminal domain gives resolvable ν_+_, ν_-_, ν_0_ and ν_dq_ transitions characteristic of ^14^N-His coupling, where the intensity ratio of the DQ to the NQI transitions increases depending on the number of coordinating ^14^N-His residues.^38,66^ The NQI transitions are broad compared to those obtained previously,^23,25,26,41^ likely reflecting heterogeneous couplings from a low affinity coordination environment determined previously in NMR experiments.^29^ We next performed ESEEM on the 1:1:1 ternary mixture of the separate N-terminal and C-terminal domains and Cu^2+^ – the resulting spectrum from this “*in trans*” experiment shows only the ESEEM transitions from the ^15^N-N-terminal domain as annotated in Figure 4C. These findings show that the N-terminal domain presents the high affinity Cu^2+^ binding segment of PrP^C^.

We next performed ESEEM and HYSCORE on both natural abundance and ^15^N-labeled full-length PrP^C^ each with one equivalent of Cu^2+^. Sensitive to the same nuclei as ESEEM, HYSCORE is a two-dimensional experiment that complements ESEEM by introducing an additional pulse in its sequence, providing peak separation and correlations between nuclear spin manifolds.^40^ The ESEEM spectrum of natural abundance PrP^C^ shows the three NQI peaks at ν_-_ = 0.73 MHz, ν_0_ = 1.34 MHz, and ν_+_ = 1.62 MHz, along with the DQ peak, ν_dq_ = 4.45 MHz (**Figure 4D**). The corresponding HYSCORE spectrum contains cross peaks that match the transitions observed by ESEEM. Of note are the cross peaks at (4.0 MHz, 0.7 MHz) and (4.0 MHz, 1.5 MHz) which are consistent with (ν_dq_, ν_-_) and (ν_dq_, ν_+_), respectively. We also observe the diagonally symmetric cross peaks representing (ν_-_, ν_dq_) and (ν_+_, ν_dq_). The ESEEM spectrum of ^15^N-PrP^C^ gives the four energy transitions: ν_α_ = 0.27 MHz, ν_β_ = 2.72 MHz, ν_nz_ = 1.50 MHz, and 2ν_α_ = 0.67 MHz (**Figure 4E**). The corresponding ^15^N-HYSCORE spectrum contains cross peaks connecting manifolds associated with the transitions observed by ESEEM. Of note are the prominent cross peaks at (0.4 MHz, 2.7 MHz) and the weaker cross peaks at (0.8 MHz, 2.7 MHz), which are consistent with (ν_α_, ν_β_) and (2ν_α_, ν_β_), respectively.

With the ESEEM and HYSCORE spectra of the individual peptide domains and full-length PrP^C^, we are now positioned to analyze LigPrP^C^ with one equivalent of Cu^2+^. The ESEEM spectrum reveals energy transitions from Cu^2+^ coordinating simultaneously to ^14^N-His and ^15^N-His residues (**Figure 4F**). For example, there is a clear DQ peak from ^14^N-His coupling and a HF peak (e.g., ν_α_) from ^15^N-His coupling. These findings demonstrate that the Cu^2+^ coordination environment in LigPrP^C^ involves His residues from both the N-terminal and C-terminal domains. The ESEEM spectrum of LigPrP^C^ potentially has significant overlap of ^14^N and ^15^N peaks at frequencies below 1.5 MHz. To thoroughly resolve the spectral details, we obtained the HYSCORE spectrum of LigPrP^C^, which reveals the intense ^15^N cross peaks associated with (ν_α_, ν_β_) and (2ν_α_, ν_β_). Additionally, the (ν_dq_, ν_+_) cross peak derived from ^14^N-His coupling is clearly resolvable, but with an intensity that is significantly weaker than that observed for unlabeled, full-length PrP^C^. Taken together, these findings support the assignment of coordination from two or more OR His residues, along with coordination from C-terminal His residues.

Given the demonstrated involvement of at least one C-terminal His residue, we next applied site-directed mutagenesis to LigPrP^C^ to test for participation in the coordination environment of His139 and His176 (**Figure 1C**). We mutated the C-terminal His residues to Tyr since its side chain has similar steric properties to that of His but does not coordinate to Cu^2+^.^29,67^ Experiments were performed with both of the single His→Tyr mutants and the double mutant LigPrP^C^(H139Y/H176Y). The relevant PrP^C^ C-terminal domains were prepared and ligated to ^15^N-N-terminal domain, purified and characterized using the same protocols described earlier. ESEEM of LigPrP^C^(H139Y/H176Y) with one equivalent of Cu^2+^ gives a DQ peak reduced to less than 5% observed for LigPrP^C^, confirming that either His139 or His176, or both, contribute to the coordination environment (**Figure 5**). We note that the very weak residual DQ peak for the double mutant and ^15^N-N-terminal PrP^C^ may arise from the ammonium chloride reagent, which is ∼99% ^15^N (1% ^14^N), used to prepare this segment. In the full-length constructs, there may also be couplings to ^14^N elsewhere in the protein C-terminal domain.

**Figure 5.**
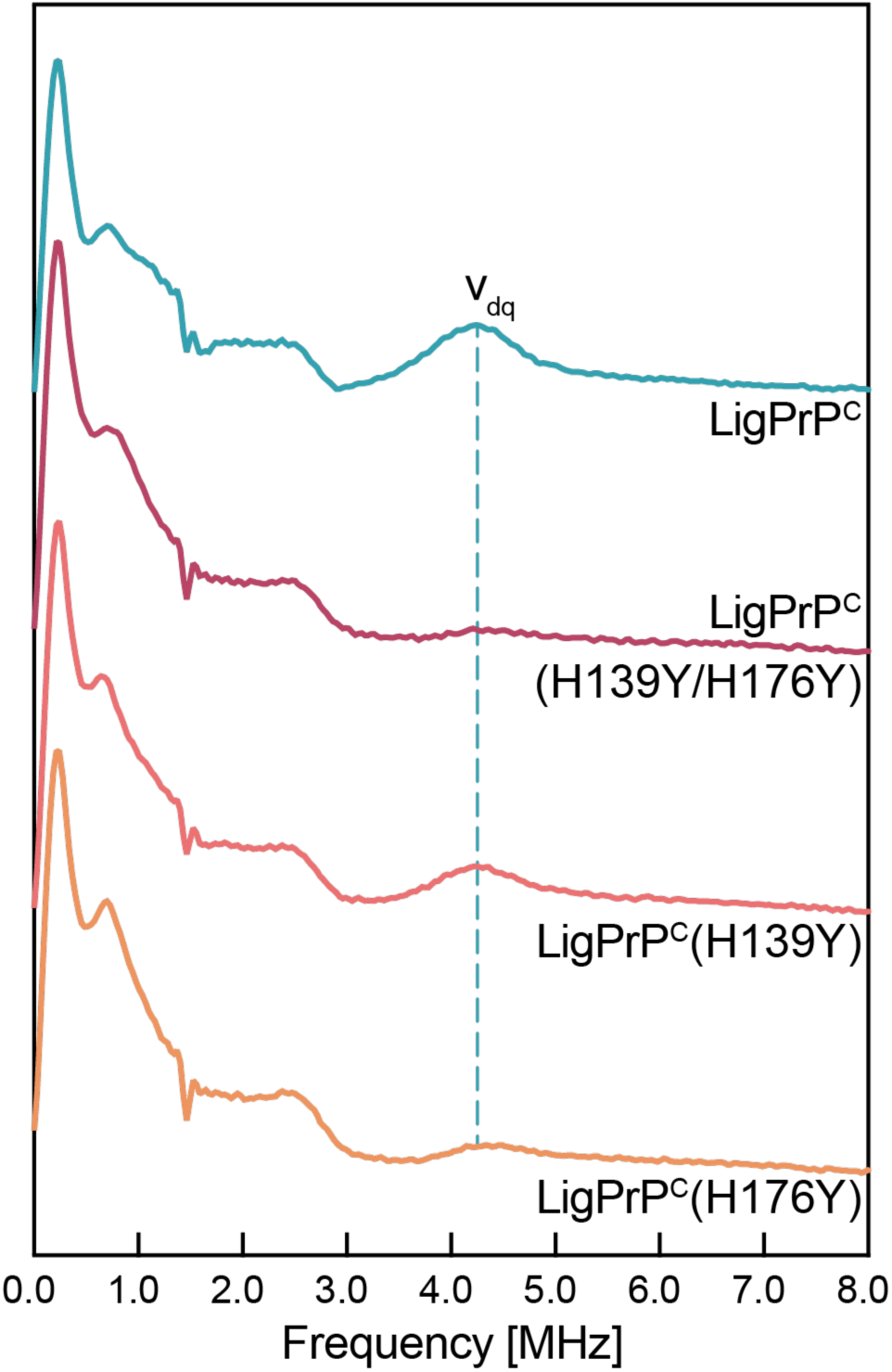
The Intensity of the DQ peak in the ESEEM of LigPrP^C^ Mutants as a Measure of C-terminal His Coordination. All spectra give energy transitions associated with ^15^N-coupling, including the 2ν_α_ ∼ 0.7 MHz, indicating multiple OR His coordination. The DQ peak from ^14^N coupling (vertical cyan dashed line at ∼ 4.3 MHz) is observed in the ESEEM spectra of LigPrP^C^ (cyan) and LigPrP^C^(H139Y) (salmon). The spectrum of LigPrP^C^(H139Y/H176Y) (dark magenta) has a DQ peak of less than 5% of the DQ peak of LigPrP^C^ when referenced against ^15^N-N-terminus alone (Figure 4C). Similarly, the intensity of the DQ peak of LigPrP^C^(H176Y) is significantly weaker than LigPrP^C^ or LigPrP^C^(H139Y) at approximately 36% of that observed for the latter.

Next, each C-terminal His was individually mutated, producing fusion products referred to as LigPrP^C^(H139Y) and LigPrP^C^(H176Y). Unlike LigPrP^C^(H139Y/H176Y), the ESEEM spectrum of LigPrP^C^(H139Y) gives a clear DQ peak of intensity similar to that of LigPrP^C^, suggesting that His139 does not contribute to copper coordination (**Figure 5**). Conversely, the ESEEM spectrum of LigPrP^C^(H176Y) is similar to that of LigPrP^C^(H139Y/H176Y), with a DQ peak that is significantly weaker than observed for LigPrP^C^. Comparison of these two ESEEM spectra suggests that His176 is the primary contributor responsible for the DQ signal observed for LigPrP^C^. However, the slight intensity of the DQ peak observed for LigPrP^C^(H176Y) suggests that there may be an equilibrium between coordination environments with one favoring His139 and the other His176, but with His176 being the dominant state. It is unlikely that both His139 and His176 simultaneously coordinate to the copper center given the distance between their β carbons of >20 Å. (Additional modeling allowing for His side chain rotamers finds that the distance of closest approach between imidazole nitrogen atoms of His139 and His176 is ∼16 Å (**Figure S3**).)

This work combined protein ligation and segmental isotope labeling with advanced EPR to provide rigorous details of copper coordination in PrP^C^. Unlike our previous efforts, this work directly tested for C-terminal His coordination. Both the ESEEM and HYSCORE spectra of LigPrP^C^ reveal the 2ν_α_ peak arising from ^15^N-N-terminal coordination, consistent with participation of at least two OR His residues. Moreover, the DQ peak arising from ^14^N-His coordination, from the globular C-terminal domain, is persistent in both LigPrP^C^ and LigPrP^C^(H139Y), demonstrating that His176 also coordinates to the copper center. However, when compared to spectra obtained from the separate domains combined with Cu^2+^ *in trans*, only ^15^N signals are observed. Consequently, formation of the copper complex stabilizing the *cis* interaction requires the intervening segment linking the two domains. Previous fitting and analysis of the ^15^N-PrP^C^ CW-EPR, which resolved superhyperfine features in the perpendicular region, demonstrated 4N coordination.^29^ Taken together, our data support a model of the *cis* interaction wherein three OR His residues and one C-terminal His, His176, coordinate to Cu^2+^, forming a square planar complex.

The *cis* interaction is a crucial regulatory mechanism, facilitated by Cu^2+^, for inhibiting the intrinsic neurotoxicity of PrP^C^. Our previous work identified the C-terminal interface responsible for the *cis* interaction to the diffuse acidic patch on its surface, which contains two highly conserved His residues, His139 and His176.^29^ Our results determine that not only does Cu^2+^ simultaneously coordinate to His residues in both domains but further reveals that His176 is the primary contact point stabilizing the *cis* interaction (**Figure 6**). To further explore the molecular details of the domain-domain interface, we carried out simulations with AlphaFold 3.^68^ Simulations were performed with the separate N-terminal and C-terminal sequences along with one Cu^2+^ ion. (We note that interacting domains must be added as separate sequences.) Several high probability structures were predicted, one of which identified a coordination environment composed of three OR His residues and His176, consistent with our experimental findings (**Figure 6**). Finally, we note that His176 is rigorously conserved, as is the composition of the OR domain, in mammalian sequences, further demonstrating the contributions of these features to the structure and stability of PrP^C^.^69^

**Figure 6.**
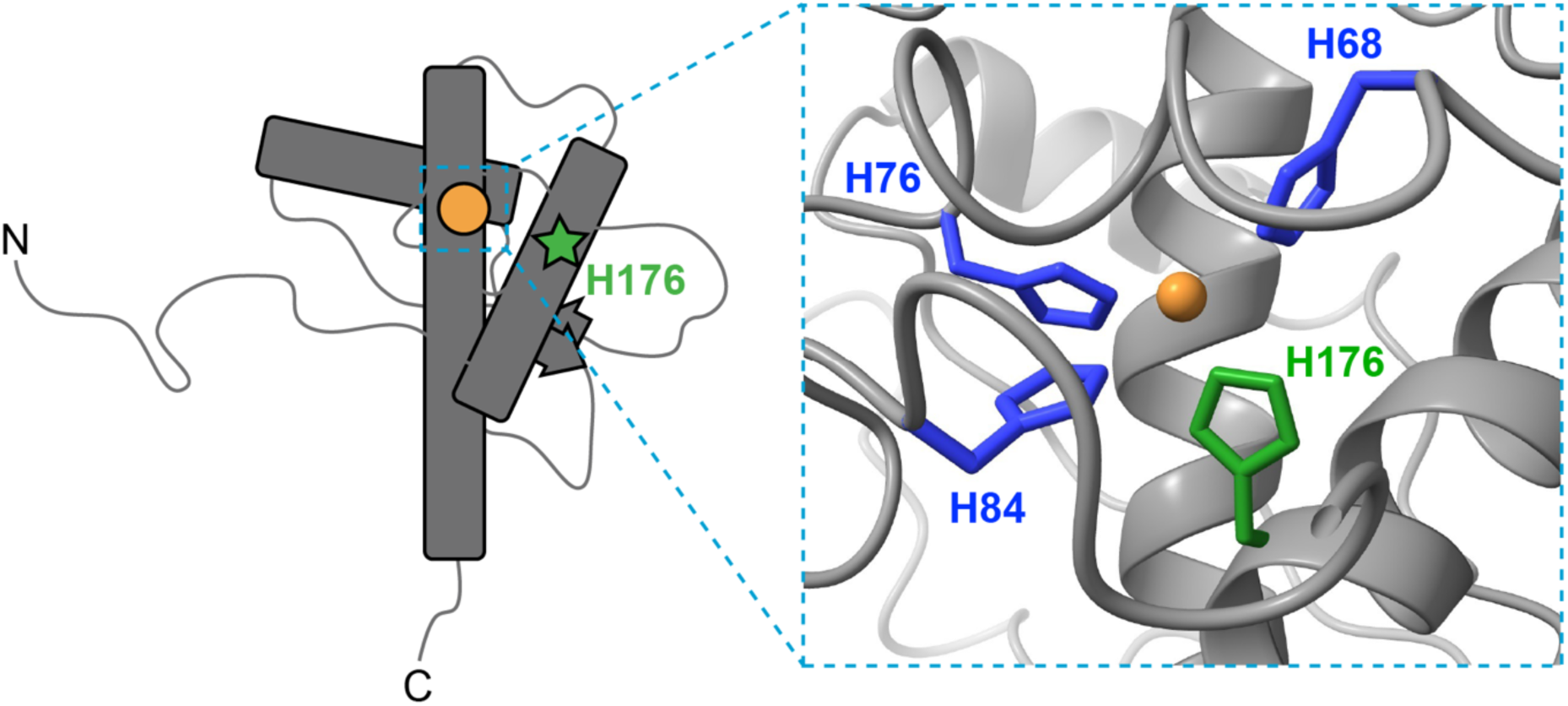
An AlphaFold 3 Model of the *Cis* Interaction. AlphaFold 3 simulations were performed with the polypeptide strands PrP^C^(23-117) and PrP^C^(118-230), with one equivalent of Cu^2+^. The graphic (right) shows the result of one high probability simulation. Simultaneous coordination by three OR His residues (blue) and C-terminal His176 is consistent with our EPR findings.

## Methods

### Protein Constructs, Expression and Purification for the Cellular Prion Protein (PrP^C^)

All recombinant prion protein (PrP) constructs coded for mouse (*Mus musculus*), mPrP. Full-length recombinant mPrP, mPrP(23-230) was in the pJ414 vector (DNA 2.0). The N-terminal construct with the sortase recognition site (LPETGG), mPrP(23-117)-LPETGG, was in the pD454-SR vector (ATUM). The C-terminal construct with a Tobacco Etch Virus (TEV)-cleavable N-terminal 6xHis tag, His-mPrP(118-230), was in the pHis::Parallel1 pET22b(+) vector, a customized, generated in-house based off the Parallel vector series kindly gifted by the Partch Lab at UCSC. Gibson cloning was utilized to generate this C-terminal construct. The first set of primers (Invitrogen) linearized the pHis::Parallel1 pET22b(+) vector, removing the N-terminal vector artifact (GAMDPEF) which usually remains after TEV cleavage, using Phusion® High-Fidelity PCR Master Mix (New England Biolabs). Once linearized, the pHis::Parallel1 pET22b(+) vector was extracted from a 1% agarose gel using the GeneJET Gel Extraction Kit (Thermo Fisher Scientific). The second set of primers (Invitrogen) amplified the mPrP sequence, coding for residues 118-230 in the pJ414 vector and added flanking regions allowing the amplified mPrP sequence to ligate with the linearized pHis::Parallel1 pET22b(+) vector. Finally, a Gibson reaction cloned the amplified mPrP(118-230) sequence into the pHis::Parallel1 pET22b(+) vector using Gibson Assembly Master Mix (New England Biolabs). This construct, pHis::Parallel1 pET22b(+)-mPrP(118-230), was then transformed into *E.coli* DH5α(DE3) (Invitrogen), extracted using MiniPrep kits (Qiagen), and validated by DNA sequencing conducted at the UC Berkeley DNA Sequencing Facility.

All PrP^C^ constructs were expressed in *E.coli* BL21(DE3) (Invitrogen). Bacteria expressing either mPrP(23-230) or His-mPrP(118-230) grew in Luria Broth (LB) media (Research Product International). Bacteria expressing mPrP(23-117)-LPETGG grew in M9 minimal media supplemented with 99% isotopically enriched ^15^N-ammonium chloride (^15^NH_4_Cl) at 1 g/L (Cambridge Isotopes). All cells grew at 37 °C until reaching an optical density (OD) of ∼1.0 at 600 nm, and then expression was induced with 1 mM isopropyl-1-thio-D-galactopyranoside (IPTG). All the cells grew for an additional 16 hours at 37 °C, except the cells expressing mPrP(23-230) which grew at 25 °C.

All recombinant mPrP constructs were extracted from inclusion bodies using 8 M guanidium chloride (GdnHCl) pH 8 at room temperature. Once extracted, only mPrP(23-230) was purified by Ni^2+^-immobilized metal ion chromatography (IMAC) Fast Protein Liquid Chromatography (FPLC) (AKTA) and eluted in 5 M GdnHCl pH 4.5. Then, its native disulfide bond formed over two days at 4 °C by bringing up the pH to 8 using KOH. mPrP(23-230) was then desalted into 10 mM potassium acetate (KOAc) pH 4.5 and purified by reverse-phase High Pressure Liquid Chromatography (HPLC) (Shimadzu) on a C4 prep column (Grace). Once ^15^N-mPrP(23-117)-LPETGG was extracted, it was immediately dialyzed into 10 mM KOAc pH 4.5 and purified by reverse-phase HPLC on a C4 prep column (Grace). His-mPrP(118-230) was also oxidatively refolded and desalted like mPrP(23-230) after extraction, but then cleaved by 6xHis-TEV in TEV reaction buffer (50 mM Tris pH 7.5, 0.5 mM Ethylenediaminetetraacetic acid (EDTA) and 1 mM Dithiothreitol (DTT)) overnight at 4 °C and finally purified by reverse-phase HPLC on a C4 prep column (Grace). The purity and identity of all mPrP constructs were verified by Liquid Chromatography-Mass Spectometry (LC-MS) (Sciex). Finally, all verified mPrP constructs were lyophilized, and stored at -20 °C until use.

### Protein Constructs, Expression and Purification for Sortase A Heptamutant

The Sortase A heptamutant (SrtA 7M, EC 3.4.22.70) was cloned into pET30b vector to be expressed with solubilizing C-terminal His6X tag (a gift from the Partch Lab at UCSC). SrtA 7M-6xHis was expressed in *E.coli* BL21(DE3) (Invitrogen) and the bacteria grew in LB media (Research Product International) at 37 °C until reaching an OD ∼0.6 at 600 nm. Next, protein expression was induced with 1 mM IPTG, with cell growth continuing for an additional 16 hours at 18 °C. After cell growth, harvest, lysis and clarification of lysate, the protein was captured using Ni-NTA affinity chromatography (Qiagen). SrtA 7M-6xHis was further purified by size exclusion chromatography on a HiLoad 16/600 Superdex 75 prep grade column (Cytiva) in 50 mM Tris pH 7.5, 150 mM NaCl, and 10% glycerol. Finally, SrtA 7M-6xHis was aliquoted, flash frozen and stored at -70 °C until use.

### Segmental labeling of PrP^C^

Sortase-mediated ligation (SML) reactions created the segmentally isotopically labeled (SIL) mPrP. We followed the protocol for C-terminal labeling by Antos et al. (2017), but with several adjustments. Specifically, Sortase A 7M-6xHis enzyme ligated the glycine peptide probe (mPrP(118-230)) to the uniformly ^15^N-labeled target protein (mPrP(23-117)-LPETGG). Before mixing, each reactant had to be solubilized in ultrapurified water and added 1 M KOH to increase the pH of the stock solutions to about 8. Then, each reactant stock was slightly diluted into 1X of the sortase reaction buffer (SRB, 50 mM Tris pH 7.4 and 150 mM NaCl). The reaction was run with dialysis for 30 minutes at 25 °C in SRB pH 7.4. Longer reaction times often resulted in partial protein precipitation. SIL mPrP was purified using Ni^2+^-immobilized metal ion chromatography (Ni^2+^-IMAC) on a HiTrap column (Cytiva). The purity and identity of SIL mPrP was verified by LC-MS (Sciex), lyophilized, and stored at -20 °C until use.

### Circular Dichroism Spectroscopy

Equilibrium CD spectra were measured using a CD spectrophotometer (J-1500, JASCO Inc., Easton, MD). mPrP(118-230) constructs (WT, H139Y, H176Y, and H139Y/H176Y), full-length PrP^C^ constructs (WT PrP^C^, LigPrP^C^, LigPrP^C^(H139Y), LigPrP^C^(H176Y), LigPrP^C^(H139Y/H176Y)), and the buffers were collected in the far-UV region (190-300 nm) using a 1 mm path length quartz cuvette. The C-terminal PrP^C^ constructs were measured in 1X SRB as described above and at a working concentration of 10 μM. The full-length LigPrP^C^ samples measured by CD were aliquots from the EPR samples and diluted to a final concentration of 10 μM. A total of 4 scans were accumulated for each of the proteins and the appropriate buffer CD spectrum was subtracted from each protein CD spectrum. Data were collected every 0.1 nm with a 4 nm bandwidth using a digital integration time of 4 sec and a scan speed of 50 nm/min. CD data were processed using the JASCO Spectra Manager Spectra Analysis package (JASCO Inc., Easton MD).

### Sample Preparation for Electron Paramagnetic Resonance (EPR)

Protein stocks were lyophilized protein dissolved in degassed Milli-Q®-purified water such that all proteins were allowed to fully solubilize and refold prior to all experiments. Then, the protein was diluted to a final concentration of 100 µM into 10 mM MES pH 6.1 and 20% (v/v) glycerol which served as cryoprotectant.^70^ CuCl_2_(*aq*) was added to a working concentration of 100 µM. Final sample volumes of 150 µL were loaded into quartz tube with a 3-mm I.D. and 4-mm O.D. (Wilmad). Lastly, samples were flash frozen by submerging into liquid N_2_.

### Continuous Wave (CW)-EPR

CW-EPR measurements were obtained at 121 K on an ElexSys E500 CW-EPR spectrometer (Bruker). Equipment included a temperature controller and an ER4122SHQE high resolution resonator. All experiments were operated at X-band frequency (∼9.3 GHz). The center field was set to 3100 G and swept for 2000 G. The modulation amplification was set to 8 Gauss and the modulation frequency was set to 100 kHz. The conversion time was 20.48 ms. The data were simulated using EasySpin 6.0.6 toolbox within the MATLAB R2024b software suite (The Mathworks Inc.).^61^

### Electron Spin Echo Envelope Modulation (ESEEM) EPR

ESEEM measurements were performed at 20 K^71^ on a ElexSys E580 X-band EPR spectrometer (Bruker). Relevant equipment included a Model 336 Temperature Cryogenic Controller (Lake Shore), Cold Edge Cryogen-free Cooling Unit (Bruker), and a MD4 resonator. All experiments were operated at X-band frequency (∼9.7 GHz). Magnetic field held constant at 3360 G, corresponding to the maximum intensity of the Cu^2+^ spectrum. The three-pulse ESEEM sequence followed: π/2 – τ – π/2 – T – π/2 – τ – echo.^36^ The parameters were π/2 = 8 ns, τ = 210 ns, T = 12 ns initially and incremented in steps of 16 ns.^41^ ESEEM spectra was collected using four-step phase cycling.^72^ The time domain signal was phased, intensity normalized, fit to an exponential decay for background subtraction, zero-filled to 2048 points, and Fourier transformed using Bruker XEPR software.

### Hyperfine Sublevel Correlation Spectroscopy (HYSCORE) EPR

HYSCORE measurements were obtained at 20 K on the same spectrometer described above. The four-pulse HYSCORE sequence used is as follows: π/2 – τ – π/2 – t_1_ – π – t_2_ – π/2 – τ – echo.^40^ The parameters were π/2 = 8 ns, π = 16 ns, t = 210 ns, T1 and T2 = 40 ns initially and incremented in steps of 16 ns. HYSCORE spectra was collected using eight-step phase cycling. The time domain signal was analyzed using Hyscorean.^73^ The time domain signals utilized Tukey apodization and were zero-filled to 1024x1024 points. Lastly, the HYSCORE spectra were obtained with a 2D Fourier transformed.

## Supporting Information

Supporting Information: SDS-Page gels from optimizing all conditions of SML, CW spectra of all constructs used in this study and additional His rotomer modeling.

## Data Availability

Raw data are available in https://github.com/Millhauser-Lab/PrP-Segmental-Labeling-Copper-EPR

## Acknowledgments

Dr. Ray Harold (UCSC) is thanked for technical assistance and mentorship for the development of the SML system. We sincerely thank Professor Carrie Partch (UCSC) for her advice and generous contribution of plasmids critical to this work. We appreciate Professor Seth Rubin for his recommendations on experimental design. The authors are grateful to Amanda Smart (UCSC) for critical reading and comments on the manuscript.

## Author Contributions

**Francesca A. Pavlovici:** Conceptualization (Lead), Methodology (Lead), Validation (Lead), Data Curation (Lead), Formal analysis (Lead), Investigation (Lead), Writing – Original Draft (Lead), Writing – Review & Editing (Lead), Visualization (Lead)**. Kevin Singewald:** Conceptualization (Supporting), Methodology (Supporting), Validation (Lead), Data Curation (Lead), Software (Lead), Formal analysis (Lead), Resources (Supporting), Writing – Original Draft (Supporting), Writing – Review & Editing (Lead), Funding acquisition (Supporting), Supervision (Supporting)**. Samuel Kaplan:** Investigation (Supporting). **Eefei Chen:** Investigation (Supporting), Resources (Supporting), Data Curation (Supporting). **Glenn L. Millhauser**: Conceptualization (Lead), Methodology (Supporting), Validation (Lead), Resources (Lead), Writing – Original Draft (Lead), Writing – Review & Editing (Lead), Supervision (Lead), Project administration (Lead), Funding acquisition (Lead)

## Funding Sources

This research was supported by the National Institutes of Health (R35GM131781, G.L.M) and National Institute of General Medicine Sciences Institutional Research and Academic Career Development Award (K12GM139185, K.S.). The pulsed EPR instrument was funded through NIH grant S10OD024980.

## For Table of Contents Only

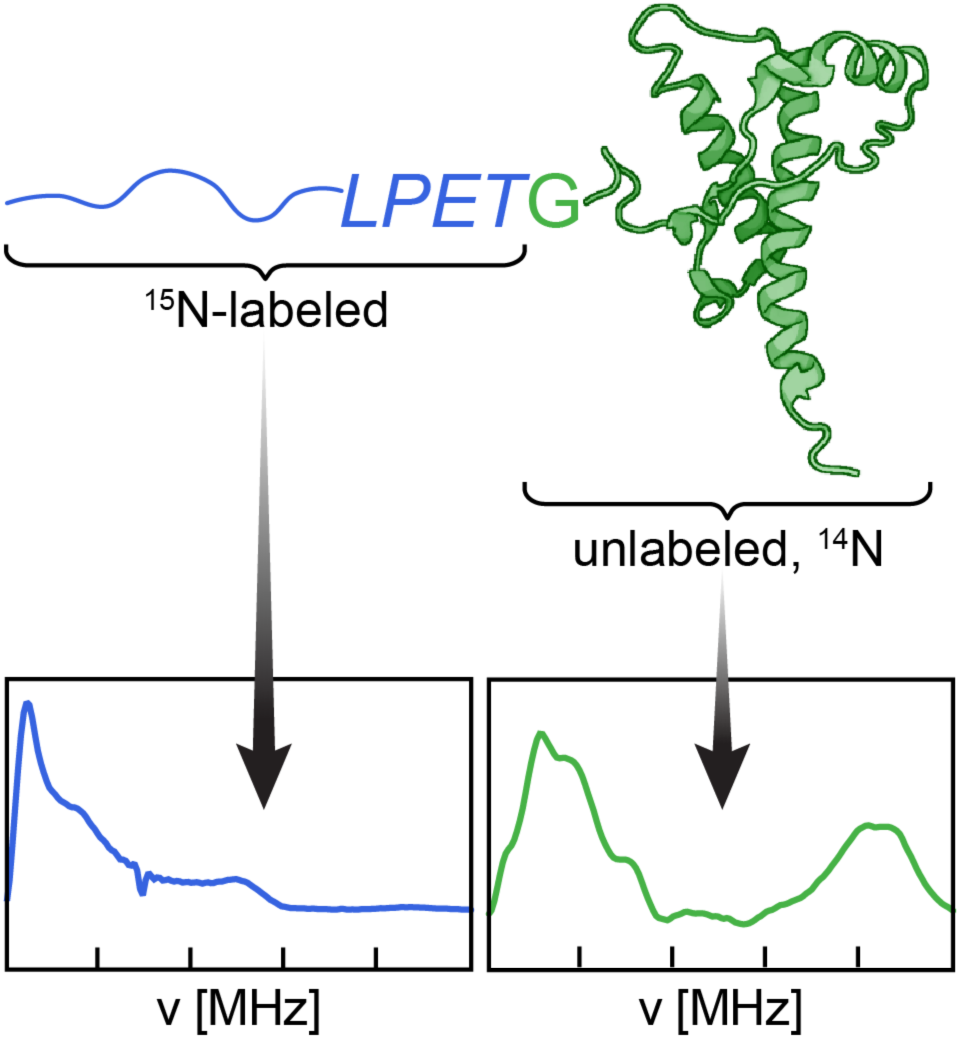

## Notes

https://github.com/Millhauser-Lab/PrP-Segmental-Labeling-Copper-EPR

### Competing Interest Statement

The authors have declared no competing interest.

### Summary of Updates

This version contains corrections and additional referencing.

