## Supporting Information for "Segmental Isotope Labelling of the Prion Protein: Identification of a Key Residue for Copper-Mediated Interdomain Structure"

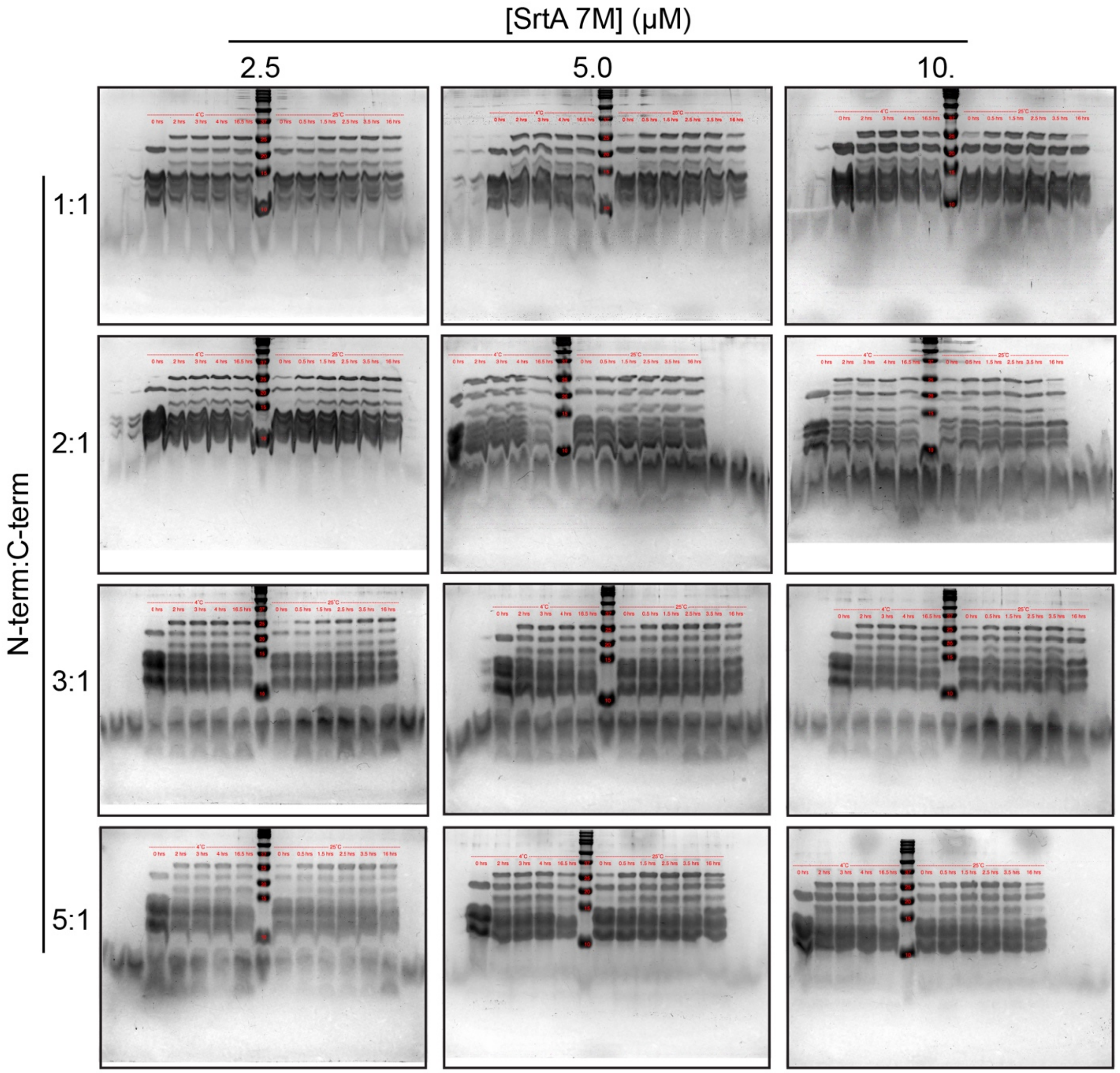


**Figure S1. The conditions of ligations are tested and visualized by SDS-PAGE gel.** Each reaction was 100 μL. Time points were taken over the course of 16-17 hours depending on temperature. Reactions at room temperature (bands to the right of the MW ladder) reached equilibrium faster than at 4 ºC (bands to the left of the MW ladder). At both temperatures, proteins immediately crashed out at higher concentrations of N-terminal domain. Proteins remained visually soluble at lower concentrations at both temperatures. The top protein band (~25 kDa), representing the fusion product, appears the most intense across all temperature and concentration conditions at 10. μM SrtA 7M. Taken together, 30 mins at room temperature with 10:5:1 ratio is the most ideal condition to run the ligation.

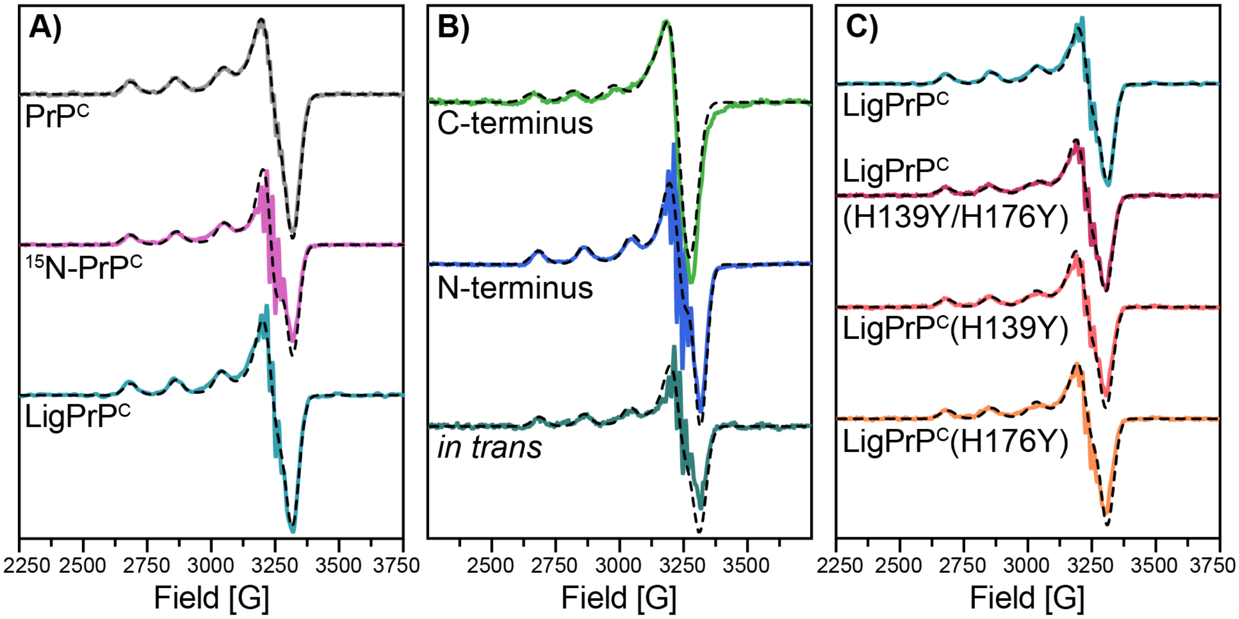


**Figure S2. Fitted CW EPR spectra of protein constructs with one equivalent of $Cu^{2+}$ used for this study. A)** The $g_{||}$ and $A_{||}$ values derived from fitting the CW spectra of PrPC (gray), $^{15}$N-PrP$^{C}$ (pink) and LigPrP$^{C}$ (cyan) align with our previously published values of Component 3 $Cu^{2+}$ binding. The CW spectrum of $^{15}$N-PrPC has sharp superhyperfine peaks, typical of $Cu^{2+}$ and $^{15}$N-nuclei coupling. **B)** The fitted CW spectrum of the C-terminus alone (green) does not result in $g_{||}$ and $A_{||}$ values associated with Component 3 $Cu^{2+}$ binding. The CW spectra of the $^{15}$N-Nterminal domain alone (blue) and the domains *in trans* (dark cyan) suggest Component 3 $Cu^{2+}$ binding based on its $g_{||}$ and $A_{||}$ values. The middle and bottom spectra give sharp superhyperfine peaks, indicative of $^{15}$N-coupling. **C)** Like LigPrP$^{C}$, the fitted CW spectrum of LigPrP$^{C}$(H139Y) produces $g_{||}$ and $A_{||}$ values that define Component 3 $Cu^{2+}$ binding. Like the C-terminal domain alone, the CW spectra of LigPrP$^{C}$(H139Y/H176Y) and LigPrP$^{C}$(H176Y) are found to not coordinate to $Cu^{2+}$ in Component 3 binding. The superhyperfine peaks are stronger for all LigPrP$^{C}$, but not as defined as homogenous $^{15}$N-protein constructs, suggesting that not all the ligands coordinating to $Cu^{2+}$ are labeled.

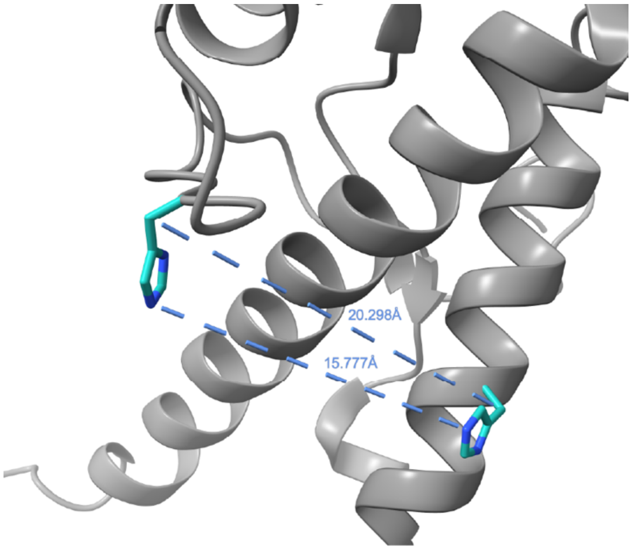


**Figure S3. Modeling the distance between His139 and His176.** Representation of mouse PrP$^{C}$ C-terminal fragment (mPrP(121-231)), PDBID: 1XYX (grey). The graphic zooms in to His139 (cyan) and His176 (cyan). The distance between the β carbons of His139 and His176 is over 20 Å. With rotation of the Cα-Cβ bond, the closest distance between the imidazole nitrogen atoms (blue) is approximately 16 Å.
